# Characterization of Congestive Heart Failure Phenotype in a Murine Model of JAK2V617F-positive Myeloproliferative Neoplasm

**DOI:** 10.64898/2026.09.24.754163

**Authors:** Xiaochuan Sun, Haoyi Zheng, Huichun Zhan

## Abstract

**Background:** Murine models are widely used to investigate cardiovascular disease (CVD), yet many established models rely on invasive surgery or genetic manipulation to induce hyperlipidemia and therefore may not fully capture the gradual cardiovascular consequences of mutant hematopoiesis. Tie2-Cre^+^/^−^FF1^+^/^−^ (Tie2FF1) mice express JAK2V617F mutation in both hematopoietic and endothelial cells and develop spontaneous CVD on a standard chow diet.

**Methods:** We applied a comprehensive, non-invasive cardiovascular phenotyping protocol to 1-year-old Tie2FF1 mice and age-matched controls. Assessments included transthoracic echocardiography, coronary flow reserve by Doppler ultrasound, quantitative analysis of coronary arteriolar structure and perivascular fibrosis, endocardial histology and subendocardial collagen deposition, and histologic evaluation of pulmonary and hepatic congestion.

**Results:** Tie2FF1 mice demonstrated left ventricular dilation, increased cardiac mass, and mildly reduced systolic function without obstructive epicardial coronary disease. Coronary microvascular remodeling was evident by arteriolar narrowing and increased perivascular fibrosis, while coronary flow reserve was markedly reduced in an exploratory cohort. Endocardial endothelial disruption and increased subendocardial fibrosis were also observed. In addition, Tie2FF1 mice showed increased lung and liver weights, pulmonary fluid accumulation, alveolar septal thickening, pulmonary vascular fibrosis, hepatic sinusoidal dilatation, and centrilobular hepatocyte swelling, providing objective evidence of clinically relevant heart failure.

**Significance:** These findings establish Tie2FF1 mice as a spontaneous, non-invasive model with features of coronary microvascular and endocardial endothelial dysfunction and heart failure with mildly reduced left ventricular ejection fraction. The integrated approach described here provides a practical framework for comprehensive murine cardiovascular phenotyping beyond traditional atherosclerosis- and surgery-based models.

## Introduction

Clonal hematopoiesis of indeterminate potential (CHIP) is defined by the presence of somatic mutations in at least 2% of circulating white blood cells in the absence of overt hematologic disease[1]. Its prevalence increases with age, affecting approximately 10–20% of individuals older than 70 years[2-5]. Although CHIP confers only a modest annual risk of hematologic malignancy (approximately 0.5–1%), it is now recognized as an important cardiovascular risk factor, associated with a 2-to 4-fold increased risk of cardiovascular disease (CVD) and worse clinical outcomes[2-6]. The magnitude of this risk is comparable to, and in some settings may exceed, that associated with traditional cardiovascular risk factors such as diabetes, hypertension, hyperlipidemia, and smoking.

Since the 1970s, murine models have become essential tools for cardiovascular disease (CVD) research because of their rapid breeding, genetic tractability, and relatively low cost, enabling mechanistic and therapeutic studies that are impractical in larger animal models. Unlike humans, in whom plasma cholesterol is transported predominantly by low-density lipoprotein (LDL), wild-type mice carry most circulating cholesterol in high-density lipoprotein (HDL) and are therefore relatively resistant to atherosclerosis[7]. To overcome this limitation, hyperlipidemic Ldlr^−^/^−^ (LDL receptor–deficient) mice are widely used; when fed a Western/high-fat diet, they develop marked hypercholesterolemia and accelerated atherosclerosis because of impaired LDL clearance. In this setting, transplantation of mutant hematopoietic cells has shown that clonal hematopoiesis accelerates atherosclerotic plaque progression via macrophage-driven inflammation[6, 8, 9]. These studies establish a strong link between clonal hematopoiesis and accelerated atherosclerosis. However, homozygous LDL receptor deficiency is a rare genetic disorder that causes severe familial hypercholesterolemia[10], limiting the broader clinical relevance of this model. Moreover, Ldlr^−^/^−^-based models primarily recapitulate large-vessel premature or early onset atherosclerosis and do not adequately capture other important manifestations of CVD, including coronary microvascular dysfunction, cardiomyopathy, and heart failure. Surgical ligation or constriction of the coronary artery or aorta has also been used to generate ischemic heart failure or pressure-overload cardiomyopathy and demonstrated that clonal hematopoiesis promotes increased myocardial inflammation and adverse remodeling[11, 12]. Although these surgical models of CVD are powerful in establishing causality, they rely on severe, artificial surgical injuries which are far more abrupt and severe than the chronic cardiovascular stress experienced by most patients with CHIP and may obscure the nuances of CHIP-mediated chronic effects. In addition, coronary ligation and aortic constriction are technically demanding with substantial inter-operator variability, contributing to variability in the severity of cardiac remodeling and dysfunction.

Among CHIP-associated mutations, the acquired kinase mutation JAK2V617F carries the highest cardiovascular risk: individuals with JAK2V617F mutant CHIP have up to a 12-fold increased risk of coronary heart disease and ischemic stroke[4, 6], and even very low mutant allele burdens (0.06-1.7%) are linked to more than a 5-fold increase in CVD risk[13]. JAK2V617F is detectable in approximately 0.2% of middle-aged individuals[14] and is the principal driver mutation in myeloproliferative neoplasms (MPNs), clonal stem cell disorders characterized by HSPC expansion and overproduction of mature blood cells. Notably, 40-50% of MPN patients develop arterial or venous thrombosis, with CVDs being a leading cause of morbidity and mortality in these patients[15]. Vascular endothelial cells (ECs) are central regulators of cardiovascular function. Although JAK2V617F is primarily expressed in hematopoietic cells, it has also been detected in vascular ECs from some patients[16-19]. Over the past several years, our laboratory has developed three non-invasive Ldlr^+/+^ murine models to study the contributions of mutant hematopoietic cells, mutant ECs, and their interactions to CVD[20-23]. These studies identified endothelial dysfunction as a key mediator of JAK2V617F-associated CVD. Here, we use Tie2-cre^+/-^FF1^+/-^ mice, in which JAK2V617F is expressed in both hematopoietic cells (including HSPCs) and ECs. This model recapitulates key features of human MPN and develops spontaneous dilated cardiomyopathy on a standard chow diet. Built on our experience over the past several years[20, 21, 23, 24], we describe a comprehensive framework for murine cardiovascular phenotyping in the middle-aged Tie2-cre^+/-^FF1^+/-^ mice that integrates clinical, histologic, and quantitative analysis. This approach not only refines the characterization of JAK2V617F-driven CVD but also provides a broadly applicable, non-invasive platform for evaluating cardiovascular phenotypes beyond traditional atherosclerosis-focused models.

## Materials and Methods Experimental mice

JAK2V617F Flip-Flop (FF1) mice (which carry a Cre-inducible human JAK2V617F gene driven by the human JAK2 promoter; JAX stock #037558) and Tie2-Cre mice (JAX stock #008863) were obtained from the Jackson Laboratory. FF1 mice were crossed with Tie2-Cre mice to generate mice expressing JAK2V617F specifically in all hematopoietic cells (including HSPCs) and vascular endothelial cells (ECs) (Tie2-cre^+/-^FF1^+/-^, or Tie2FF1), thereby modeling the human disease in which both hematopoietic stem cells and ECs harbor the JAK2V617F mutation. All mice were bred and maintained in a specific pathogen-free animal facility at Stony Brook University. All animal experiments were performed in accordance with protocols approved by the Institutional Animal Care and Use Committee.

### Complete blood counts

Peripheral blood was obtained from the facial vein via submandibular bleeding and collected in EDTA coated tubes. The samples were then analyzed using a Vetscan Hm5 Hematology Analyzer (Abaxis) to determine the blood count.

### Flow cytometry

Cells were counted and stained with fluorochrome-conjugated antibodies by incubating cell suspensions at 4°C for 30 minutes in the dark. Samples were analyzed using a CytoFLEX flow cytometer (Beckman Coulter, Indianapolis, IN, USA). The following antibodies were used: lineage (Lin) cocktail (CD3, B220, Gr-1, CD11b, and Ter119; BioLegend), c-Kit (clone 2B8; BioLegend), Sca-1 (clone D7; BioLegend), CD150 (clone mShad150; eBioscience, San Diego, CA, USA), and CD48 (clone HM48-1; BioLegend). Bone marrow LSK cells were defined as Lin^−^ c-Kit^+^ Sca-1^+^ cells, and hematopoietic stem cells (HSCs) were defined as Lin^−^ c-Kit^+^ Sca-1^+^ CD48^−^ CD150^+^cells. Flow cytometry data were analyzed using Kaluza software (Beckman Coulter).

### Transthoracic echocardiography

Transthoracic echocardiography was performed in mildly anesthetized, spontaneously breathing mice maintained under 1-3% isoflurane in oxygen at a flow rate of approximately 1L/min using a Vevo 3100 high-resolution imaging system (VisualSonics Inc, Toronto, Canada), as previously described. Parasternal long-axis and sequential parasternal short-axis views were acquired to assess global and regional wall motion. Left ventricular (LV) end-diastolic and end-systolic dimensions were measured at the mid-ventricular level, and LV ejection fraction and fractional shortening were calculated using standard formulas to evaluate systolic function. All cardiac structural and functional measurements were obtained while maintaining the heart rate between 400 and 500 beats per minute.

Coronary flow reserve (CFR) was measured in the septal coronary artery (SCA) using a previously established method[25]. The SCA was identified in the parasternal long-axis view using B-mode and color-Doppler imaging, with the aortic valve and pulmonary artery serving as anatomical landmarks. Pulsed-wave Doppler was performed to capture coronary blood flow with the sample volume positioned in the center of the SCA lumen, and the angle between the ultrasound beam and the direction of coronary blood flow was maintained at ≤60°. Peak diastolic coronary flow velocity was measured under baseline conditions (1% isoflurane) and during hyperemia induced by 2.5% isoflurane for 1 minute. CFR was calculated as the ratio of hyperemic to baseline peak diastolic coronary flow velocity.

### Heart histology

Hearts were excised, transected at the base from the ascending aorta, and fixed in cold 4% paraformaldehyde overnight at 4°C with gentle shaking. Following fixation, hearts were washed three times with PBS at room temperature to remove residual fixative. Paraffin-embedded sections (5μm thick) were stained with hematoxylin and eosin (H&E) or Masson’s trichrome using standard protocols and reagents from Sigma (St. Louis, MO). Whole-slide images were acquired using a NanoZoomer S20 Digital Slide Scanner (Hamamatsu Photonics K.K., Japan).

Coronary microvasculature was evaluated on H&E-stained sections using Image J (NIH) as previously described[21, 24]. The ratio of luminal radius to wall thickness was determined for individual coronary arterioles (10-100 um in diameter), averaging four different measurement (top-bottom, left-right, and both diagonals). Arterioles were considered stenotic when the luminal radius-to-wall thickness ratio was <1.

Perivascular and subendocardial collagen content was assessed on Masson’s Trichrome-stained sections, where collagen fibers appear blue, muscle fibers red, and nuclei black[24]. For perivascular collagen quantification, intramyocardial vessels with circular profiles and intact morphology (10–100 µm in diameter) were selected. The collagen-positive (blue) area surrounding each vessel was measured in ImageJ and expressed as a percentage of the total vessel area. For subendocardial collagen quantification, collagen content beneath the endocardial surface of both left and right ventricles was calculated as the ratio of collagen-positive subendocardial area to the length of the corresponding endocardial segment.

### Lung and liver histology

For lung histology[26, 27], mice were exsanguinated by retro-orbital blood collection before tissue harvest to minimize residual intravascular blood. The left lobe was carefully dissected at the hilum near the origin of the main bronchus and gently blotted on tissue paper to remove excess surface fluid. Wet left lobe lung weight was recorded immediately, after which the tissue was dried at 65°C for 36 hours and reweighed. Total lung water, used as an index of pulmonary edema, was calculated as the difference between wet and dry weights. The right cranial lung lobe was collected for histological analysis and fixed immediately in 4% paraformaldehyde overnight at 4°C with gentle rocking.

For liver analysis[28, 29], mice were similarly exsanguinated by retro-orbital blood collection before tissue harvest to minimize residual intravascular blood. The liver was carefully separated from the diaphragm and surrounding tissues, removed from the abdominal cavity, gently blotted to remove excess surface fluid, and weighed. The left liver lobe was collected for histological analysis and fixed immediately in 4% paraformaldehyde overnight at 4°C with gentle rocking.

After fixation, lung and liver tissues were washed three times with PBS to remove residual fixative, paraffin-embedded, and sectioned at 5 μm. Sections were stained with H&E or Masson’s trichrome using standard protocols and reagents from Sigma (St. Louis, MO). Whole-slide images were acquired using a NanoZoomer S20 Digital Slide Scanner (Hamamatsu Photonics K.K., Japan).

### Statistical analysis

Data are presented as the mean ± standard error of mean (SEM). Differences between two groups were assessed using an unpaired two-tailed Student’s t-test for normally distributed data or the Mann-Whitney U test for non-normally distributed data. Differences among multiple groups were assessed using ANOVA, followed by post hoc multiple-comparison tests. Figures were generated using GraphPad Prism 10 (GraphPad Software, San Diego, CA).

## Results

### Spontaneous Cardiovascular Dysfunction in Tie2FF1 mice

As we previously reported, Tie2FF1 mice develop a robust MPN phenotype[30-32]. After 20 weeks of age, they exhibited significant leukocytosis (mainly neutrophilia), thrombocytosis, splenomegaly, and HSPC expansion compared to age-matched controls (Figure 1). Because the incidence of CVD increases sharply after age 50-60 in human[33], we focused our cardiovascular assessment on middle-aged (12-month-old) mice.

**Figure 1.**
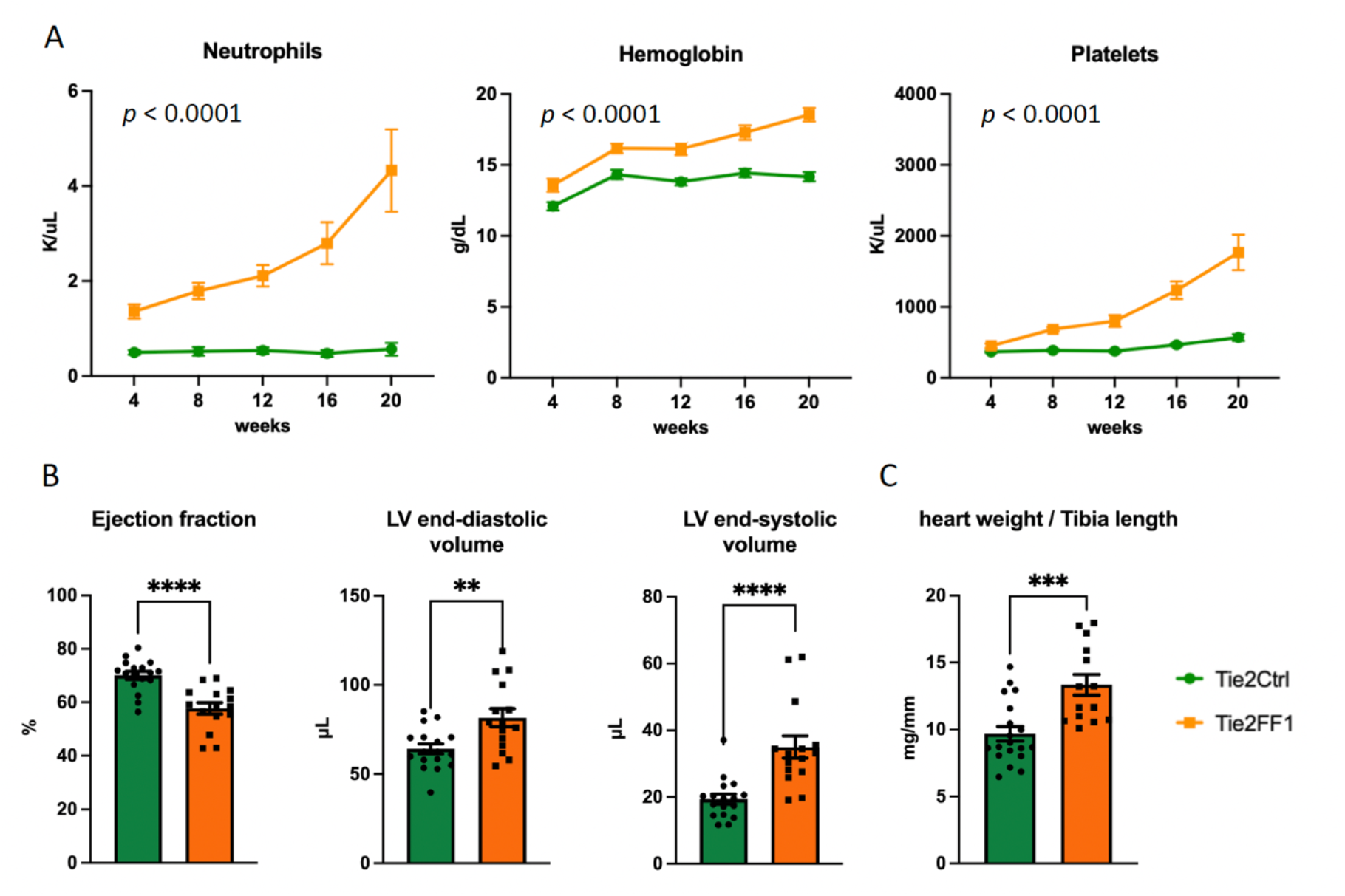
Hematopoietic and cardiovascular phenotype in Tie2FF1 mice on a regular chow diet. (**A**) Peripheral blood counts showed significantly increased neutrophil counts, hemoglobin levels, and platelet counts in Tie2FF1 mice (orange) compared with Tie2-cre control mice (green) (n = 15-17 per group). (**B**) Transthoracic echocardiography demonstrated moderately decreased LV ejection fraction with increased LV end-diastolic volume and end-systolic volumes in Tie2FF1 mice compared to age-matched Tie2-cre control mice (n = 15-17 per group). (**C**) Quantification of heart weight normalized to tibia length in 1 year old Tie2FF1 and Tie2-cre control mice (n = 14-19 per group). Statistical significance was determined by two-way ANOVA for (A) and either an unpaired Student’s t-test or Mann-Whitney U test for (B) and (C). ** p < .01; *** p < 0.001; **** p < 0.0001.

On a standard chow diet, Tie2FF1 mice developed a dilated cardiomyopathy phenotype. At 1 year of age, they had modest but significantly reduced LV ejection fraction (58% versus 70%, *p* < 0.0001), accompanied by increased LV end-diastolic volume (82 μL versus 64 μL, *p* = 0.0038) and end-systolic volume (33 μL versus 20 μL, *p* < 0.0001), compared with age-matched Tie2-Cre controls (Figure 1B). There was no echocardiographic evidence of regional wall motion abnormalities, including segmental akinesis or dyskinesis, or evidence of focal wall thinning or scarring, suggesting there was no transmural myocardial infarction. Pathologic evaluation further supported the diagnosis of dilated cardiomyopathy, with significantly increased heart size and heart weight-to-tibia length ratio in Tie2FF1 mice (Figure 1C).

### Microvascular Remodeling and Endocardial Injury in Tie2FF1 Mice

As we previously reported[32, 34], Tie2FF1 mice developed thrombosis in the right ventricle and main pulmonary arteries, while age-matched Tie2-cre control mice showed no evidence of cardiac or pulmonary thrombosis (Figure 2A). This prominent thrombotic phenotype is unique to Tie2FF1 mice, in which both hematopoietic cells and vascular ECs carry the JAK2V617F mutation, and is not observed when the mutation is restricted to either endothelial or hematopoietic cells alone[21, 22].

**Figure 2.**
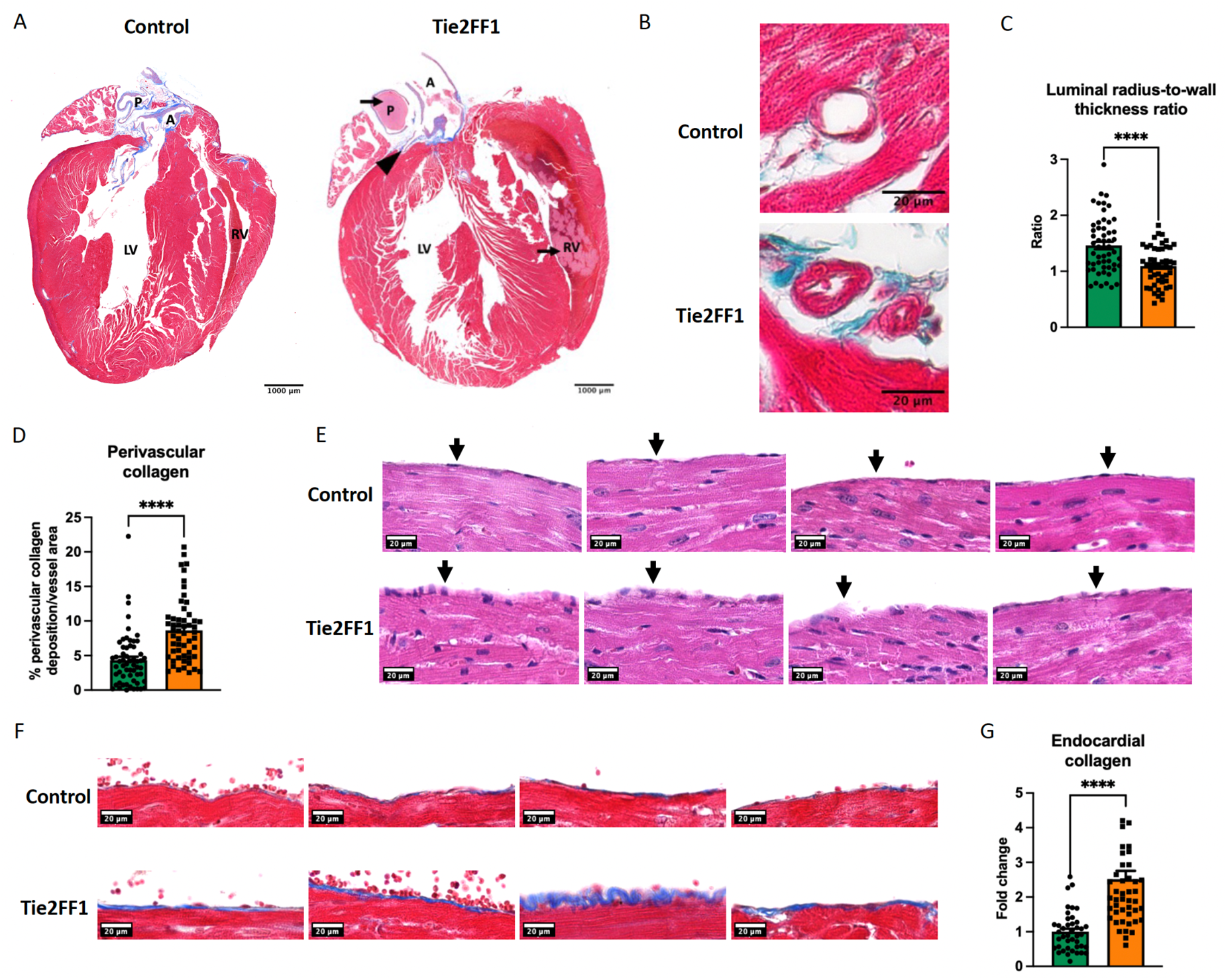
Microvascular Remodeling and Endocardial Injury in Tie2FF1 Mice. (**A**) Representative Masson’s trichrome staining of longitudinal heart sections from Tie2-cre control and Tie2FF1 mice. Note the presence of thrombus (arrow) in the right ventricle and main pulmonary artery of the Tie2FF1 mice. The epicardial coronary artery is indicated by an arrowhead. LV, left ventricle; RV, right ventricle; A, aortic root; P, main pulmonary artery. Original magnification, 20x. (**B**) Representative Masson’s trichrome staining of coronary arterioles in Tie2FF1 and control mice. (**C**) Quantification of the lumen radius-to-wall thickness ratio in non-overlapping coronary arterioles from Tie2FF1 (orange) and Tie2-cre control (green) mice (n = 4 mice per group; 50-55 microvessels analyzed per group). (**D**) Quantitation of perivascular collagen deposition around coronary arterioles in Tie2FF1 (orange) and Tie2Ctrl (green) mice (4 mice per group; 54-58 microvessels analyzed per group). (**E**) Representative H&E staining of the endocardium (arrows) in Tie2-cre control (top) and Tie2FF1 (bottom) mice. (**F-G**) Representative Masson’s trichrome staining (F) and quantitation (G) of endocardial collagen deposition across non-overlapping endocardium segments in Tie2FF1 and control mice (4 mice per group; 42-47 segments analyzed per group, average segment length 55um). Statistical significance was determined using an unpaired Student’s t-test. **** p < 0.0001.

Coronary vascular analysis revealed marked arteriole remodeling in Tie2FF1 mice, characterized by significant lumen narrowing, as reflected by a decreased luminal radius-to-wall thickness ratio (1.09 vs. 1.46, *p* < 0.0001) and a significantly increase in perivascular collagen deposition (2.2-fold, *p* < 0.0001) compared to controls (Figure 2B-D). No gross epicardial coronary atherosclerosis or myocardium infarction was detected.

The endocardium is the innermost layer of the heart chambers and consists of a specialized monolayer of ECs that play critical roles in coronary development, cardiac repair, and regeneration[35, 36] during both embryonic[37] and postnatal life[38]. Our recent work demonstrated that endocardial ECs undergo marked transcriptional reprogramming and structural disruption in a murine CVD model driven by JAK2V617F-mutant hematopoiesis and cardiometabolic stress[22]. Similar endocardial abnormalities were observed in Tie2FF1 mice, characterized by irregular lining, prominent nuclei, cellular enlargement, or increased cellularity, indicating endocardial EC activation and proliferation (Figure 2E). Masson’ trichrome staining further demonstrated a significant increase in subendocardium collagen deposition in Tie2FF1 mice (2.2-fold, *p* < 0.0001) compared with controls (Figure 2F-G).

Collectively, these findings indicate that JAK2V617-mutant hematopoiesis, together with mutant ECs, drives a non-atherosclerotic CVD phenotype characterized by microvascular dysfunction, endocardial endothelial injury, increased LV mass, and modestly reduced, but relatively preserved LV ejection fraction in 1-year old Tie2FF1 mice. This phenotype closely resembles clinical syndromes such as ischemia with non-obstructive coronary arteries (INOCA) and heart failure with preserved ejection fraction (HFpEF), in which coronary microvascular dysfunction is increasingly recognized as a key pathogenic mechanism[39, 40].

### Cardiac Microvascular Dysfunction in Tie2FF1 Mice

To further assess cardiac microvascular function, we measured coronary flow reserve (CFR) noninvasively using a Vevo 3100 high-resolution ultrasound system equipped with an MX250 (15-30MHz) transducer (Figure 3A-B). Briefly, pulsed-wave Doppler measurements were obtained from the proximal left coronary artery under baseline conditions (1.0% isoflurane) and during hyperemia induced by 2.5% isoflurane. CFR was calculated as the ratio of peak hyperemic to baseline coronary flow velocity. Because coronary flow resistance is determined primarily by the microvascular circulation, a reduced CFR (< 2.0) in the absence of epicardial coronary artery disease is a well-established indicator of coronary microvascular dysfunction. In a small exploratory cohort (n = 2-3 mice per group), Tie2FF1 mice showed a clear trend toward decreased CFR compared with controls (mean CFR, 1.06 vs. 2.25) (Figure 3C-D). Together with the marked arteriole narrowing and perivascular fibrosis observed in these mice, these findings provide evidence of both cardiac microvascular remodeling and functional impairment in the Tie2FF1 mice.

**Figure 3.**
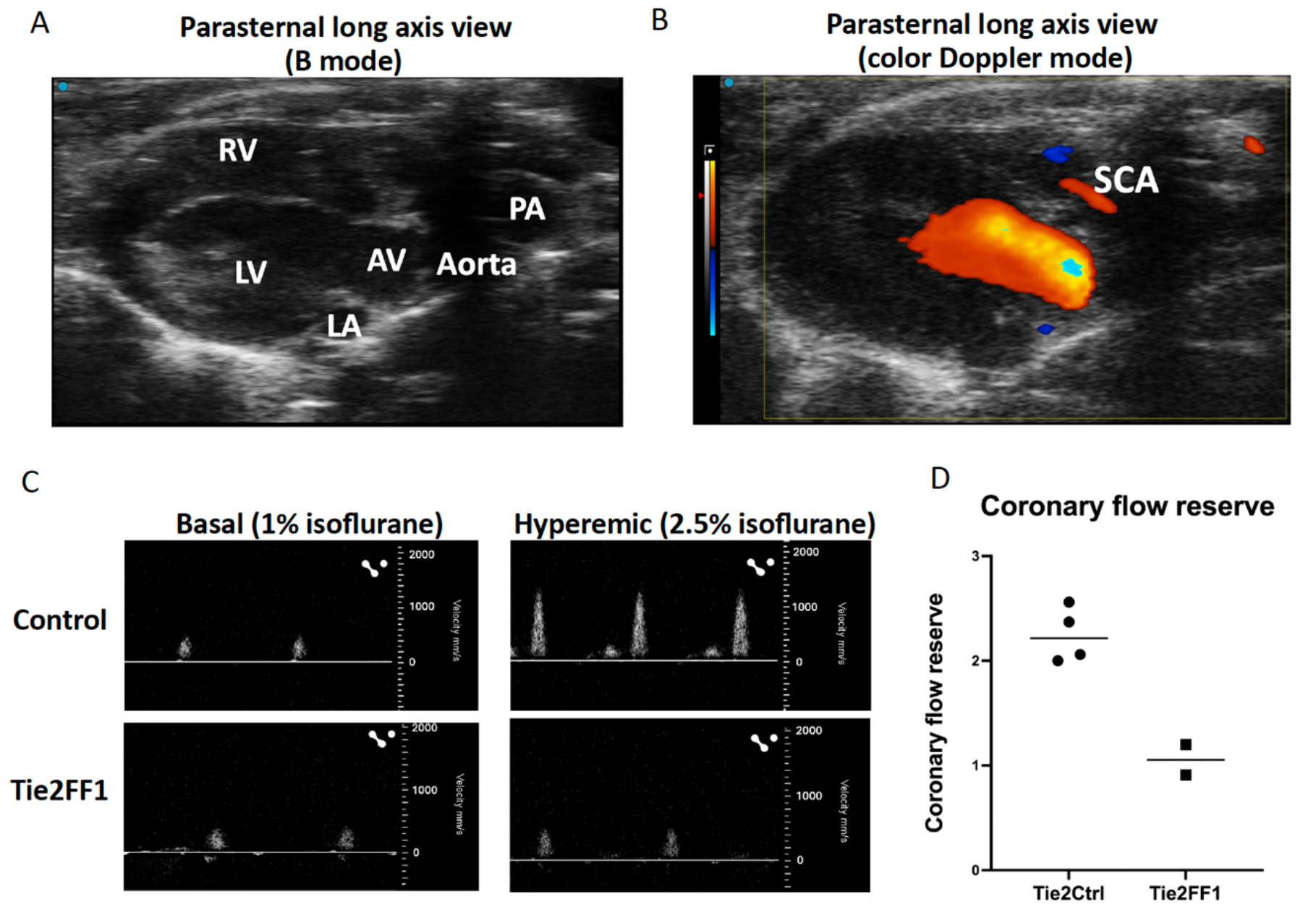
Cardiac Microvascular Dysfunction in Tie2FF1 Mice. (**A-B**) Parasternal long-axis views in B mode (A) and color Doppler mode (B) demonstrating localization of the septal coronary artery. RV: right ventricle; LV: left ventricle; LA: left atrium; PA: pulmonary artery; AV: aortic valve; SCA: septal coronary artery. (**C-D**) Representative pulsed-wave Doppler recordings of SCA flow velocity under basal conditions (1% isoflurane) and during hyperemia induced by 2.5% isoflurane (C) and CFR measurements (D) in Tie2FF1 (n=2) and control (n=4) mice.

### Pulmonary and Hepatic Congestion in Tie2FF1 Mice

Heart failure is a clinical syndrome characterized by impaired cardiac function and fluid accumulation in the lungs and peripheral tissues. Although dyspnea and fatigue cannot be assessed in mice, tissue congestion provides an objective surrogate for congestive heart failure. We therefore evaluated pulmonary[27] and hepatic[28, 29] congestion in Tie2FF1 mice.

Compared with age-matched Tie2-cre controls, Tie2FF1 mice exhibited a significant increase in left lung lobe wet weight (1.6-fold, *p* = 0.0087, n = 6-8 per group) (Figure 4A). A significantly greater fluid accumulation was also observed in the left lung lobe, assessed by the wet-dry weight difference (1.6-fold, *p* = 0.0030, n = 6-8 per group), consistent with pulmonary congestion (Figure 4B). Histologic examination further demonstrated alveolar septal thickening and increased cellularity in the lung parenchyma of Tie2FF1 mice (Figure 4C-D). In addition, Masson’s Trichrome staining revealed increased perivascular fibrosis compared with controls (Figure 4E).

**Figure 4.**
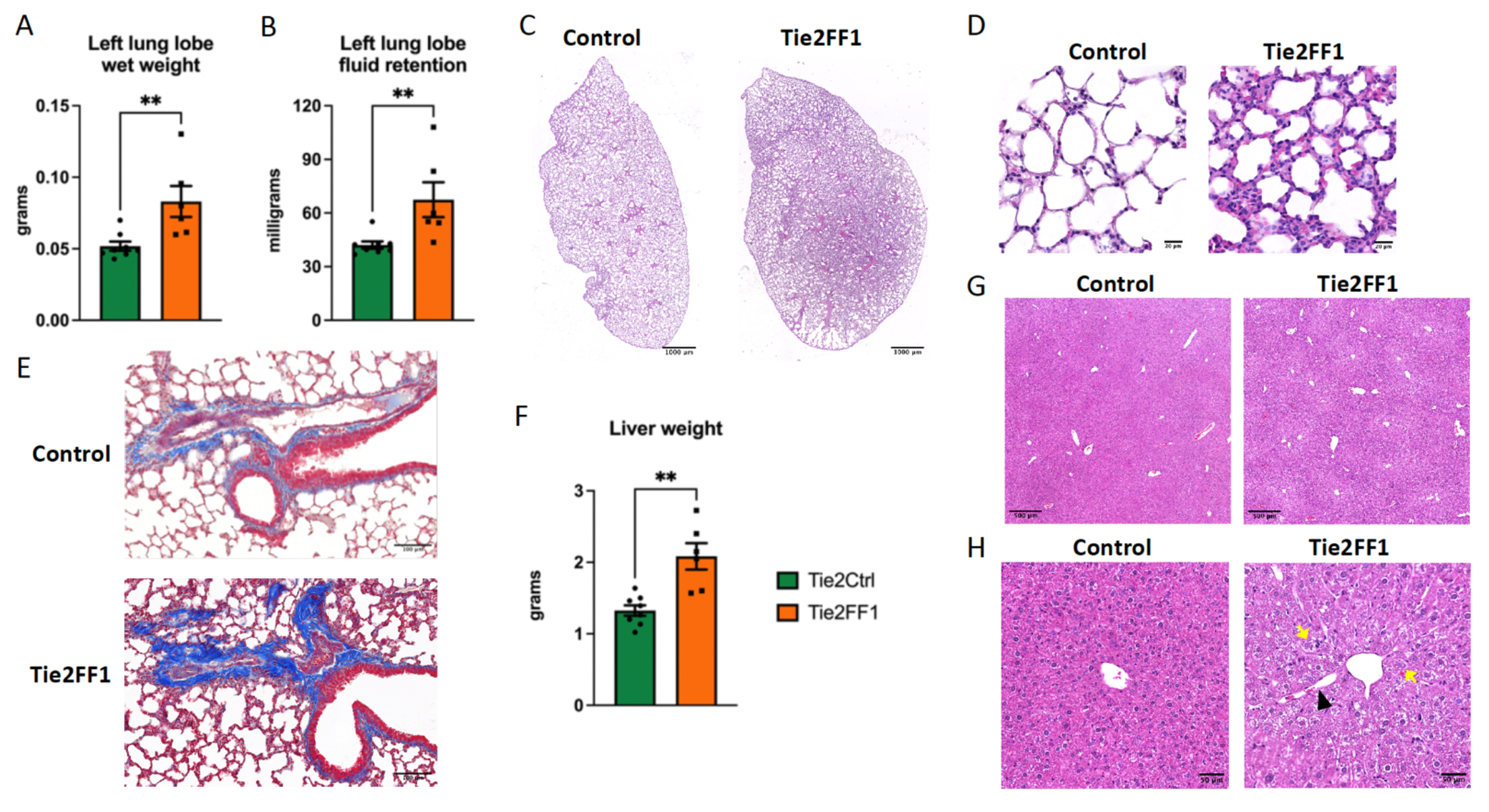
Pulmonary and Hepatic Congestion in Tie2FF1 Mice. (**A**) Left lung lobe wet weight in Tie2FF1 (orange) and Tie2-cre control mice (green) (n = 6-8 per group). (**B**) Left lung lobe fluid content, calculated as wet weight minus dry weight, in Tie2FF1 (orange) and control (green) mice (n = 6-8 per group). (**C-D**) Representative H&E-stained sections of the right upper lung lobe shown as the full field of view (C) and an enlarged view of a selected region (D), demonstrating alveolar septal thickening and increased cellularity in Tie2FF1 mice. (**E**) Representative Masson’s trichrome staining of lung sections showing increased perivascular fibrosis in Tie2FF1 mice compared to control mice. (**F**) Liver weight in Tie2FF1 (orange) and Tie2-cre control (green) mice (n = 6-8 per group). (**G-H**) Representative H&E-stained sections of the left liver lobe as the full field of view (G) and an enlarged view of a selected region (H), showing sinusoidal dilatation (black arrowhead) and centrilobular hepatocyte swelling (yellow arrow) in Tie2FF1 mice. Statistical significance was determined by an unpaired Student’s t-test for normally distributed data or the Mann-Whitney U test for non-normally distributed data. ** p < 0.01.

The liver is another major organ affected by passive venous congestion in heart failure, often manifesting as congestive hepatopathy[28, 29]. Tie2FF1 mice exhibited a significant increase in liver weight compared to controls (1.6-fold, *p* = 0.0012, n = 6-8 per group) (Figure 4F). Hematoxylin and eosin staining demonstrated classic features of congestive hepatopathy, including sinusoidal dilatation, accompanied by centrilobular hepatocyte swelling (Figure 4G-H).

Together, these findings establish objective tissue-based markers for systemic congestion and support a congestive heart failure phenotype in Tie2FF1 mice.

## Discussion

Heart failure, a progressive clinical syndrome characterized by impaired cardiac contraction or relaxation, remains a leading cause of morbidity and mortality worldwide. Although we first reported the Tie2FF1 model in 2020[20], our understanding of JAK2V617F-associated cardiovascular phenotype has evolved substantially over the past several years[20-23, 32]. Initial studies established that combined expression of JAK2V617F in hematopoietic cells and endothelial cells produces an MPN-like phenotype accompanied by spontaneous cardiovascular disease. In this study, we characterized the cardiovascular phenotype of the Tie2FF1 mice at 1 year of age using a broader cardiovascular evaluation and revealed several previously unrecognized features. In addition to LV dilation and mildly reduced ejection fraction, Tie2FF1 mice exhibited coronary arteriolar narrowing, increased perivascular fibrosis, impaired coronary flow reserve, endocardial endothelial disruption, and increased subendocardial collagen deposition.

In addition, we identified objective systemic evidence of congestive heart failure that was not fully appreciated in the original characterization of this model[20]. Tie2FF1 mice demonstrated increased lung wet weight and pulmonary fluid accumulation, alveolar septal thickening, pulmonary vascular and perivascular fibrosis, increased liver weight, sinusoidal dilatation, and centrilobular hepatocyte swelling. These findings provide tissue-level evidence of pulmonary and hepatic congestion and support the presence of clinically relevant heart failure rather than isolated abnormalities in cardiac structure or function. Because symptoms such as dyspnea or fatigue cannot be recorded in mice, and gravity-dependent peripheral edema is less pronounced in quadrupeds because of the minimal hydrostatic pressure gradient between the heart and distal limbs, we used pulmonary and hepatic congestion as objective surrogates of congestive heart failure symptoms, as these manifestations correspond to clinically relevant signs and symptoms in patients with heart failure. When considered together with increased LV mass, moderately reduced LVEF, cardiac microvascular remodeling, and impaired CFR, the phenotype shares important features with human cardiac microvascular dysfunction and HFpEF or INOCA. Importantly, these structural and functional abnormalities occurred in the absence of gross epicardial coronary atherosclerosis or myocardial infarction and without chronic cardiometabolic stress.

A major strength of this model is that cardiovascular disease develops spontaneously without surgical intervention or genetic manipulation of the LDL receptor pathway. Many murine studies of CHIP-associated cardiovascular disease have relied on either Ldlr^−^/^−^ mice fed with a Western diet to accelerate atherosclerosis[6, 8, 9] or invasive procedures such as coronary ligation and transverse aortic constriction to induce myocardial injury or pressure overload[11, 12]. These approaches have provided important mechanistic insights but also impose severe, artificial cardiovascular stress and may obscure more chronic effects of mutant hematopoiesis. In contrast, Tie2FF1 mice are maintained on an Ldlr^+^/^+^ background and develop cardiovascular abnormalities on a standard chow diet without coronary ligation, aortic constriction, or other surgical injury. This allows the cardiovascular consequences of the JAK2V617F mutation to evolve over time and provides an opportunity to study pathogenesis of CVD. Our findings expand the spectrum of JAK2V617F-associated cardiovascular disease beyond thrombosis[20, 41, 42] and atherosclerosis[9, 43] to include cardiac microvascular and endocardial endothelial dysfunction. Endocardial ECs is critical for coronary vasculature formation, cardiac repair, and regeneration[35, 36]. Together with our recent findings in mice with JAK2V617F-mutant hematopoiesis[22], these results suggest that endocardial ECs may have important contribution to JAK2V617F-associated CVDs. The findings may have important clinical implications in INOCA and HFpEF. INOCA refers to a clinical condition characterized by myocardial ischemia in the absence of significant coronary artery stenosis. Coronary microvascular dysfunction and endothelial dysfunction are the major findings in patients with INOCA[44]. HFpEF accounts for more than half of heart failure cases with poorly understood pathogenesis and limited effective therapies[45, 46]. Cardiac microvascular dysfunction, driven by systemic inflammation and endothelial dysfunction, are central mechanisms underlying HFpEF[47]. Emerging evidence associates CHIP with non-obstructive coronary artery disease, coronary microvascular dysfunction, and increased HFpEF risk[48-50]. Our models[20-23, 32], including the Tie2FF1 mice, provide a unique, noninvasive platform for investigating JAK2V617F-associated cardiac microvascular dysfunction and HFpEF. The Tie2FF1 model is not a perfect replica of HFpEF, particularly because prominent thrombosis appears to be a distinct feature of combined mutant hematopoiesis and endothelial JAK2V617F expression, which is not observed when the mutation is restricted to either endothelial or hematopoietic cells alone[21, 22].

The present study also underscores the value of comprehensive phenotyping when evaluating murine cardiovascular disease. Histological analysis or echocardiographic measurements alone may identify LV remodeling and systolic or diastolic dysfunction but may not adequately capture the broader disease process. By integrating transthoracic echocardiography, noninvasive measurement of coronary flow reserve, quantitative analysis of coronary arteriolar remodeling and fibrosis, assessment of endocardial structure, and histologic evaluation of pulmonary and hepatic congestion, we were able to define a much more complete phenotype of heart failure and cardiac microvascular disease. These observations also illustrate an important broader principle in experimental cardiovascular research: well-established animal models may yield substantially new biological information when revisited with improved technologies and a wider set of clinically relevant endpoints. The Tie2FF1 model was originally developed to investigate the interaction between mutant hematopoietic and endothelial cells in MPN-associated vascular disease. Six years later, expanded cardiovascular phenotyping revealed that this model also recapitulates features of coronary microvascular disease, endocardial endothelial injury, and congestive heart failure that were not fully characterized in the initial study. This evolution in understanding highlights the importance of longitudinally re-evaluating disease models as new questions and methodological approaches emerge.

In summary, our findings establish aged Tie2FF1 mice as a spontaneous, non-invasive model of JAK2V617F-associated cardiovascular disease on an Ldlr^+^/^+^ background. We hope that both the methodological approaches used here and the cardiovascular features described will provide a practical framework for comprehensive murine cardiovascular phenotyping and facilitate future studies examining how mutant hematopoiesis contributes to the development and progression of CVD.

## ACKNOWLEDGMENTS

This work was supported by the U.S. Department of Veterans Affairs Merit Award BX003947 and BX005584 (H.Z.) and NIH R01 CA266294 (H.Z.).

## AUTHOR CONTRIBUTIONS

X.S. performed all experiments, analyzed the data, and prepared the figures. H. Zheng provided scientific and technical expertise in cardiovascular functional assessments. H. Zhan conceived and supervised the study, interpreted the results. X.S. and H. Zhan drafted the manuscript. H. Zheng critically revised the manuscript. All authors reviewed and approved the final manuscript.

